# PINK1 supports colorectal cancer growth by regulating the labile iron pool

**DOI:** 10.1101/2022.09.27.509713

**Authors:** Brandon Chen, Nupur K. Das, Indrani Talukdar, Rashi Singhal, Cristina Castillo, Anthony Andren, Joseph D. Mancias, Costas A. Lyssiotis, Yatrik M. Shah

## Abstract

Mitophagy is a cargo-specific autophagic process that recycles damaged mitochondria to promote mitochondrial turnover. PTEN-induced putative kinase 1 (PINK1) mediates the canonical mitophagic pathway. We show that PINK1 expression is positively correlated with decreased colon cancer survival, and mitophagy is required for colon cancer growth following nutrient stress. However, the mechanism by which PINK1 maintains colon cancer growth remains equivocal. Inducible knockdown (KD) of PINK1 in a panel of colon cancer cell lines inhibited colon cancer cell proliferation, whereas disruption of other mitophagy receptors did not similarly impact cellular proliferation. Mechanistically, we observed a decrease in mitochondrial respiration, membrane hyperpolarization, accumulation of mitochondrial DNA, and depletion of antioxidant glutathione following PINK1 KD. Mitochondria are important hubs for storing iron and synthesizing iron-dependent cofactors such as heme and iron sulfur clusters. An increase iron storage protein ferritin and a decrease labile iron pool was observed in PINK1 KD cells. However, neither total cellular iron nor markers of iron starvation/overload were affected. Cellular iron storage and the labile iron pool are maintained via autophagic degradation of ferritin (ferritinophagy). Overexpressing nuclear receptor coactivator 4 (NCOA4), a key adaptor for ferritinophagy, rescued cell growth and the labile iron pool in PINK1 KD cells. We demonstrate that PINK1 regulates intracellular iron availability by integrating mitophagy to ferritinophagy. In conclusion, these results indicate that PINK1 is essential for maintaining intracellular iron homeostasis to support survival and growth in colorectal cancer cells.

## Introduction

Mitochondria are critical metabolic organelles that sustain cellular bioenergetics and biosynthetic needs^1^. The electron transport chain (ETC) integrates central carbon metabolism and redox homeostasis to support metabolic demands of the cells. To maintain a healthy mitochondrial network, organellar functions are continuously monitored via quality control mechanisms^2^. Dysfunctional mitochondria are turned over by a cargo-specific, lysosomal-dependent, autophagic degradation mechanism termed mitophagy^3^. Dysregulation of mitophagy has been associated with progression of several cancers^4^. Parkin-induced protein kinase 1 (PINK1) is a sensor of mitochondrial health, and activation of PINK1 regulates one of the most well-defined mitophagy pathways^5,6^. However, the role of PINK1 as a tumor suppressive surveillance mechanism for enhancing survival and proliferation are context dependent^78,9^.

Induction of PINK1-mediated mitophagy is triggered by loss of membrane potential from uncoupling the proton or potassium gradient^10^. Recent results illustrated that chelation of mitochondrial iron potently induced mitophagy (insert ref). The role of mitochondria in iron metabolism is well established as both iron sulfur cluster (Fe-S) and heme biosynthesis starts within this organelle. Moreover, mitochondria ETC require Fe-S cluster and heme containing protein such as cytochrome c to facilitate electron transfer. Nevertheless, the mechanistic connection between mitochondrial iron loss and mitophagy remains unclear^11,12^.

In excess, iron is cytotoxic to cells. As such, cellular iron levels are balanced by an intricate network of regulatory mechanisms. A central protein in iron handling is the iron storage protein ferritin (FTN). Preliminary data suggest most bioavailable iron uptake goes through a transient FTN intermediate to prevent oxidative damage^13^. Subsequently, FTN bound iron release is mediated by autophagic degradation of nuclear receptor coactivator 4 (NCOA4) in a process termed ferritinophagy. Following FTN degradation, lysosomal/endosomal iron can be distributed in the cell via direct lysosome-organelle contacts or chaperones. As an example, recent studies demonstrated that iron turnover from ferritinophagy is critical to support mitochondrial iron sulfur cluster biogenesis and respiration in pancreatic cancer^14,15^.

To study the connection between mitophagy and iron balance, we generated a genetic, PINK1 loss-of-function model in colorectal cancer cell models. We identified PINK1-mediated mitophagy as a critical pathway for CRC cell proliferation and mitochondrial function. The loss of PINK1 led to increased sequestration of labile iron in ferritin. The proliferative defects and cellular labile iron pool in PINK1 KD was rescued by activating ferritinophagy via nuclear receptor coactivator 4 (NCOA4) overexpression. Overall, our study suggests that disruption of the canonical mitophagy pathway contributes to cytosolic iron imbalance, which can be rescued by activating ferritinophagy.

## Results

### PINK1 is required for colorectal cancer cell proliferation *in vitro* and *in vivo*

Previously, we demonstrated that colorectal cancer cells utilize mitophagy to support metabolic rewiring under nutrient deprived conditions^16^. Mining a large colon cancer patient database demonstrated that high PINK1 expression is positively correlated with worsened progression/regression free survival (**Figure 1A**). In addition, mining the Human Protein Atlas, colorectal cancer cells robustly express PINK1 compared to several other cancers (**Figure 1B**). PINK1 is the initiating kinase that licenses phospho ubiquitin (pUb)-dependent mitophagy initiation^17^, and we generated two independent doxycycline (DOX) inducible small hairpin RNA (shRNA) targeting PINK1. To test cell proliferation and viability, we counted cell number over time to account for proliferation and utilized colony formation assay to assess the ability of cells to form single unit colonies. When comparing between WT and PINK1 KD, we decided to compare DOX treatment groups of shNT vs shPINK1 instead of between isogenic shPINK1 cells with and without DOX. This minimizes any off-target effects of doxycycline. Upon PINK1 knock down (KD), we observed proliferative defects and decreased colony forming capacity in several colorectal cancer cell lines (HCT116, SW480, RKO, HT29, MC38, CT26) (**Figure 1C and D**). KD was validated by decreased PINK1 mRNA transcript (**Supplemental figure 1A**). However, disruption of PINK1-independent mitophagy executors, parkin, BNIP3, NIX, FUNDC1 did not exhibit growth defects (**Supplement figure 1B and 1C**). In addition, analysis of PINK1 KD cells in a subcutaneous xenograft model reveled decreased tumor burden with decreased tumor volume, decreased tumor weight, decreased BrdU staining, and increased TUNEL staining following doxycycline treatment (**Figure 2A, B, C, and D**). This data demonstrates that PINK1 has an essential role in mediating CRC cell growth.

**Figure 1.**
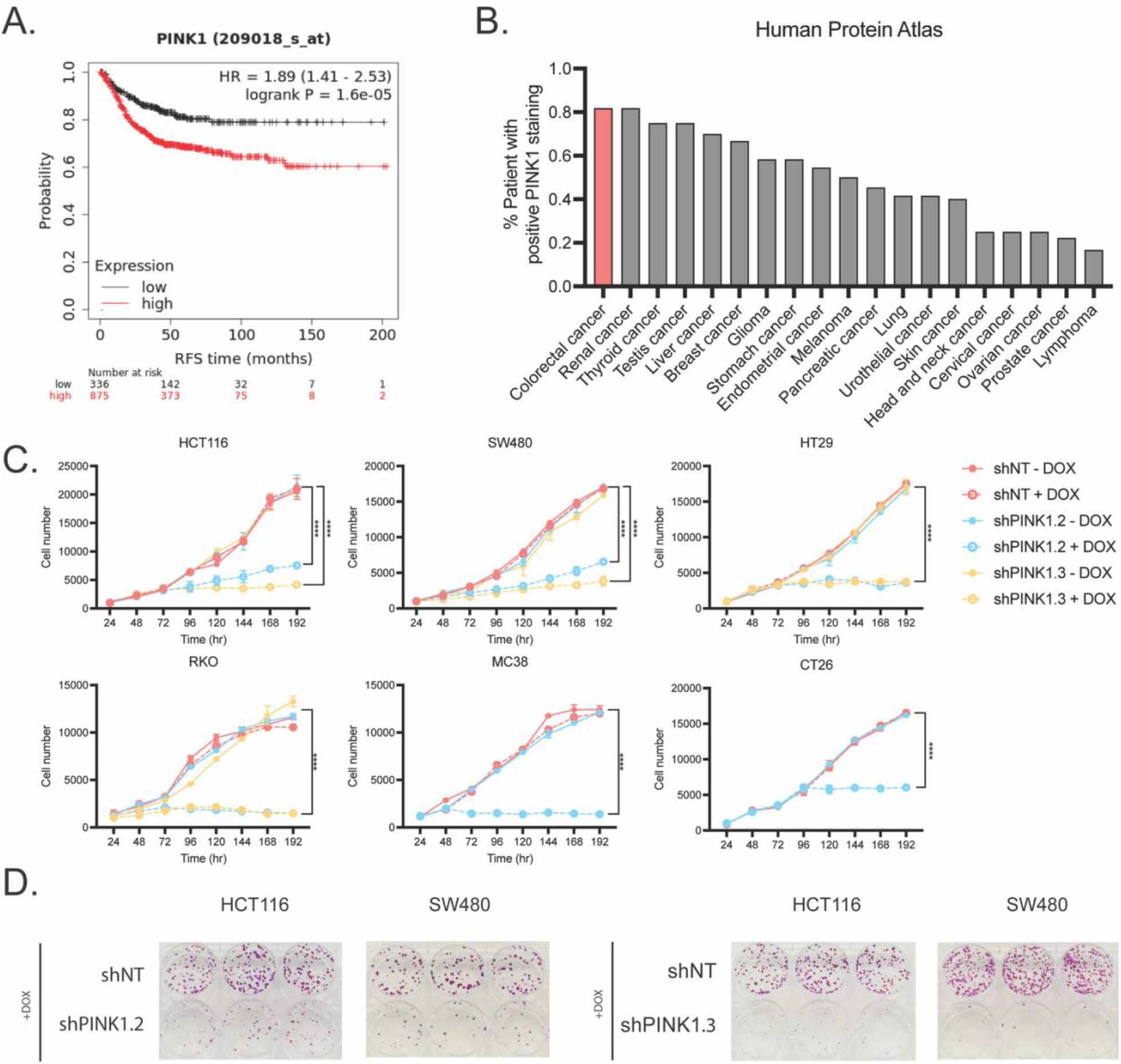
PINK1 is associated with poor patient survival and required for colorectal cancer growth in vitro. **A**. Regression free survival in colorectal cancer patients stratified on median PINK1 expression. **B**. Human Protein Atlas (HPA) data on % patient with positive PINK1 expression (Accession number HPA00193). Human colorectal cell lines (HCT116, SW480, HT29, RKO) and mouse colorectal cancer cell lines (CT26 and MC38) with doxycycline (DOX) inducible non-targeting (shNT) and 2 independent shRNA targeting human PINK1 (shPINK1.2 and shPINK1.3) or mouse PINK1 (shPINK1.2) were assessed for **C**. cell proliferation and **D**. colony formation assays following 50 ng/ml of doxycycline.

**Figure 2.**
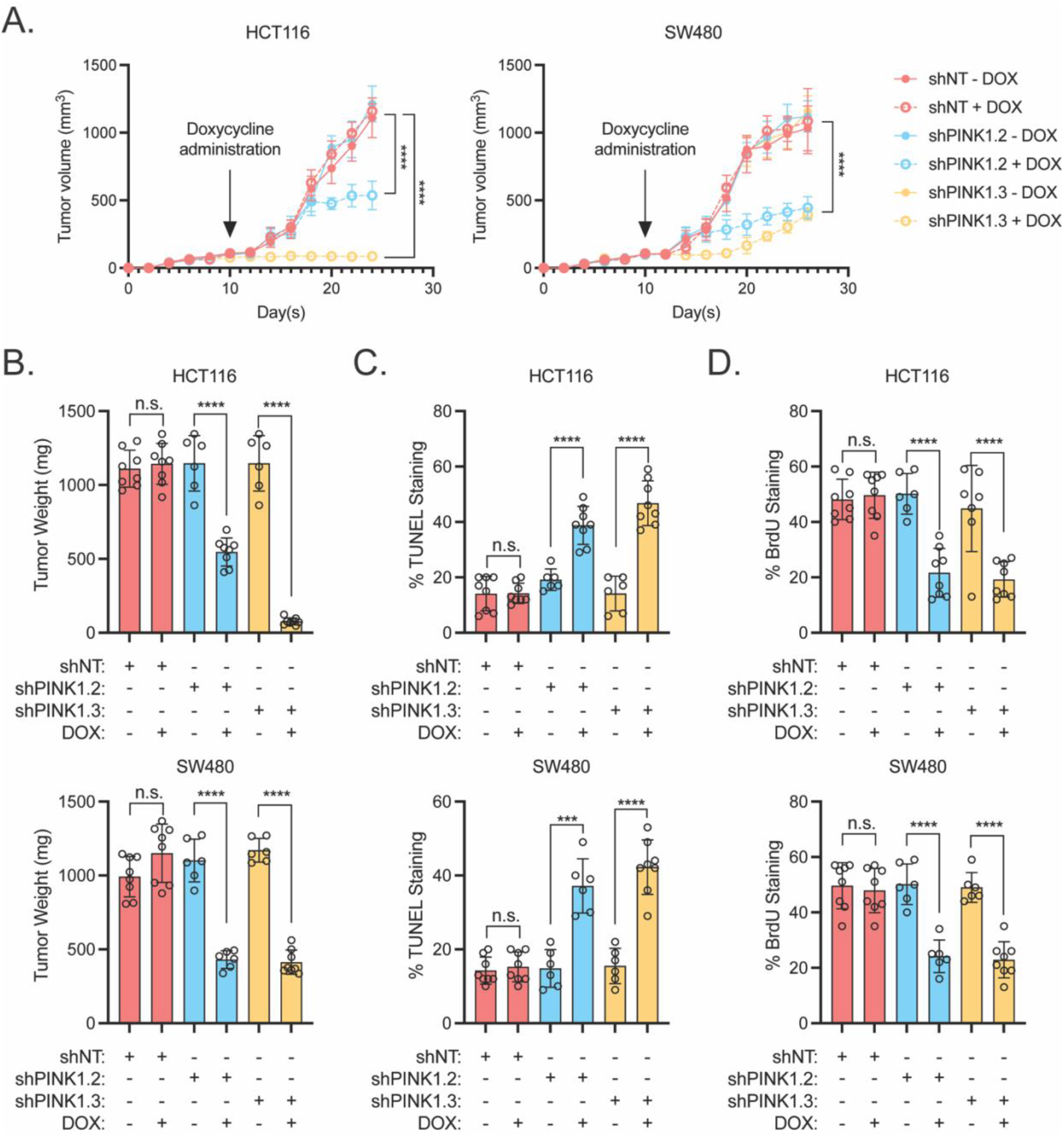
PINK1 is required for colorectal cancer in vivo growth. HCT116 and SW480 shNT, shPINK1.2, and shPINK1.3 tumors were injected into the flanks of NOD/SCID mice (N=6-8) and **A**. tumor volume, **B**. end point tumor weight, **C**. Percent (%) TUNEL staining and **D**. BrdU staining were quantified with or without doxycycline (DOX) chow. Data is presented as mean +/- the standard error of the mean. **** indicates P < 0.0001.

### Inhibition of PINK1 disrupts mitophagy and mitochondrial functions

Mitophagy is the process by which damaged mitochondria are degraded and the components are recycled to support cell growth. Upon loss of mitochondrial membrane potential, PINK1 accumulates on the outer mitochondrial membrane (OMM) and recruits E3 ligase Parkin to coordinate the decoration of OMM and OMM proteins with phospho-ubiquitin (pUb) chains^18^. pUb chains subsequently recruit autophagic adaptors such as p62 to deliver mitochondria to autophagosomes for degradation^11^. PINK1 knockdown (KD) prevented accumulation of pUb in response to carbonyl cyanide m–chlorophenylhydrazone (CCCP), a mitochondrial uncoupler (**Figure 3A**). In addition, we sought to characterize mitophagy flux utilizing a mitochondrially-targeted tandem Cox8-mCherry-GFP mitophagy reporter (**Figure 3B**). GFP is a pH-sensitive fluorophore, while mCherry is pH-insensitive. Upon engulfment of mitochondria into the acidic autophagosome, it quenches the localized GFP signal, whereas mCherry will remain fluorescent. PINK1 KD effectively decreased mitophagy flux (**Figure 3C**). Moreover, we observed increased mitochondrial DNA (mtDNA) content, which indicated accumulation of mitochondria (**Figure 3D**).

**Figure 3.**
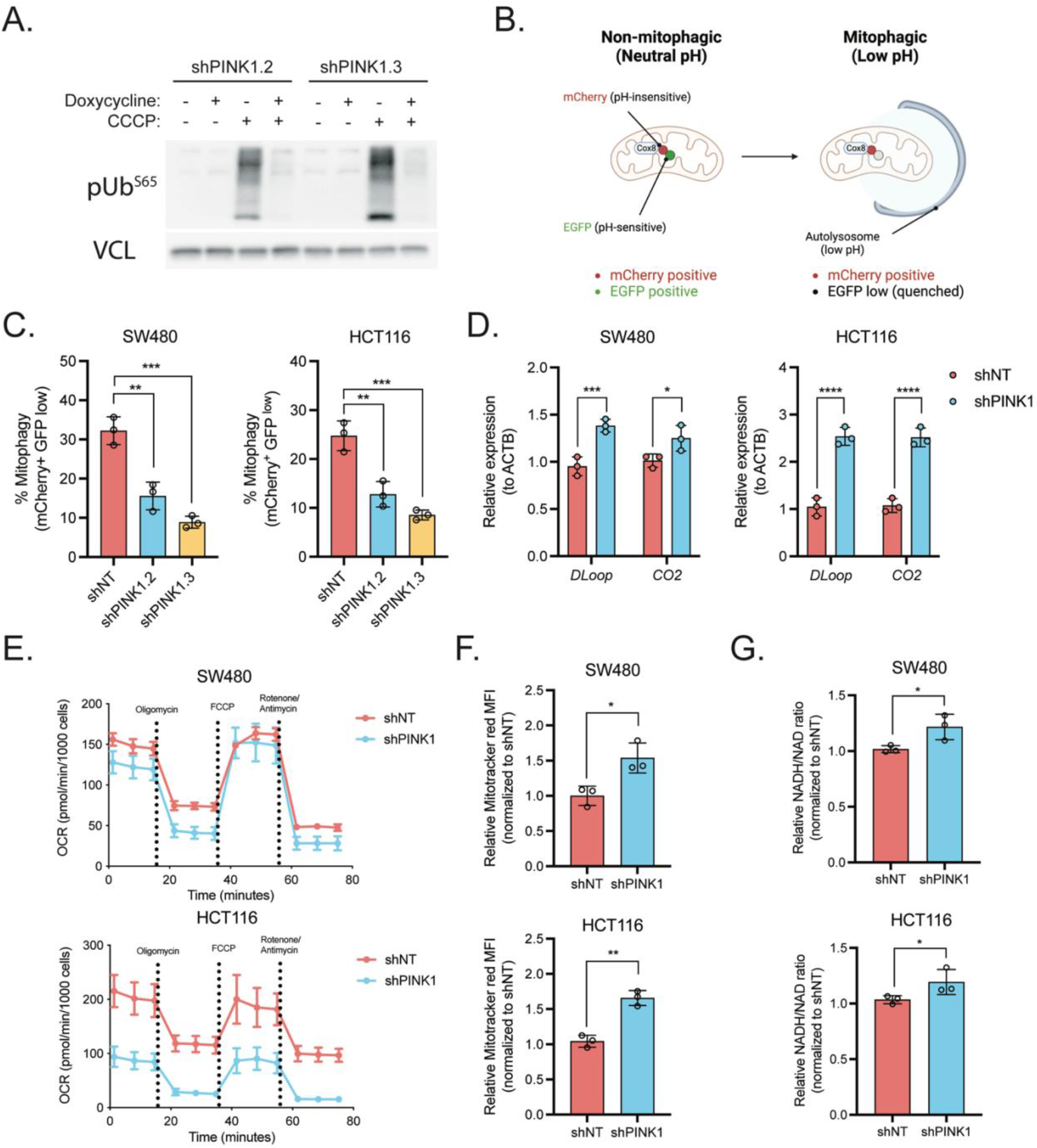
Loss of PINK1 disrupts mitophagy and contributes to mitochondrial dysfunction. **A**. Schematic of the COX8-EGFP-mCherry mitophagy reporter. **B**. Mitophagy flux was measured using the reporter. **C**. Cells were treated with vehicle or 10 μM CCCP, and phosphorylated ubiquitin (pS65-Ub) was assessed by Western blot analysis. **D**. mtDNA content was assessed using mitochondrially encoded gene D-Loop and Cytochrome c oxidase 2 (CO2). **E**. Seahorse MitoStress Test was conducted with following order and concentration of inhibitors: Oligomycin (1 μM), FCCP (1 μM), Rotenone (1 μM) and Antimycin A (1 μM). (mean -/+ SEM, N = 4 per condition). F. Mitochondrial membrane potential is measured by Mitotracker Red CMXRos. G. Relative NADH/NAD ratio. Data is presented as mean +/- the standard error of the mean, *P < 0.05, **P < 0.001, ****P < 0.0001. All experiments were done in triplicates and repeated at least three times.

Next, mitochondrial function was investigated following PINK1 KD. Seahorse analysis demonstrated lower basal oxygen consumption rate (OCR) and maximal respiration in PINK1 KD cells (**Figure 3E and supplemental figure 2A and 2B**). ETC is tightly coupled with mitochondrial membrane potential, and PINK1 KD cells had increased mitochondrial membrane potential, which is indicative of mitochondrial hyperpolarization and a potential blockade in ETC and proton motive force (**Figure 3F**). Another role of mitochondrial ETC complex I is to oxidize NADH to sustain NAD pool, which can be assessed by measuring NADH/NAD ration. We observed elevated levels NADH/NAD ration upon PINK1 KD, which contributes to cellular reductive stress (**Figure 3G**).

### Cellular redox balance and nucleotide metabolism are disrupted upon PINK1 inhibition

Mitochondria integrate the metabolism of nutrients such as glucose, glutamine, and fatty acids to coordinate energy production, the regulation of redox homeostasis, and other biosynthetic precursors. Liquid chromatography tandem mass spectrometry (LC-MS/MS)-based metabolomics was used to profile changes in central carbon metabolism upon PINK1 KD. Here, we observed a downregulation of the reduced glutathione (GSH) to oxidized glutathione (GSSG) ratio and nucleotides (i.e. ADP, AMP, UDP, IMP) in SW480 (**Figure 4A, 4B, and table 1**). Together with the bioenergetic profiling data in Figure 2, this evidence indicates that losing the ability to recycle mitochondria via PINK1-dependent mitophagy inhibits mitochondrial respiration and contributes to cellular metabolic dysfunction.

**Figure 4.**
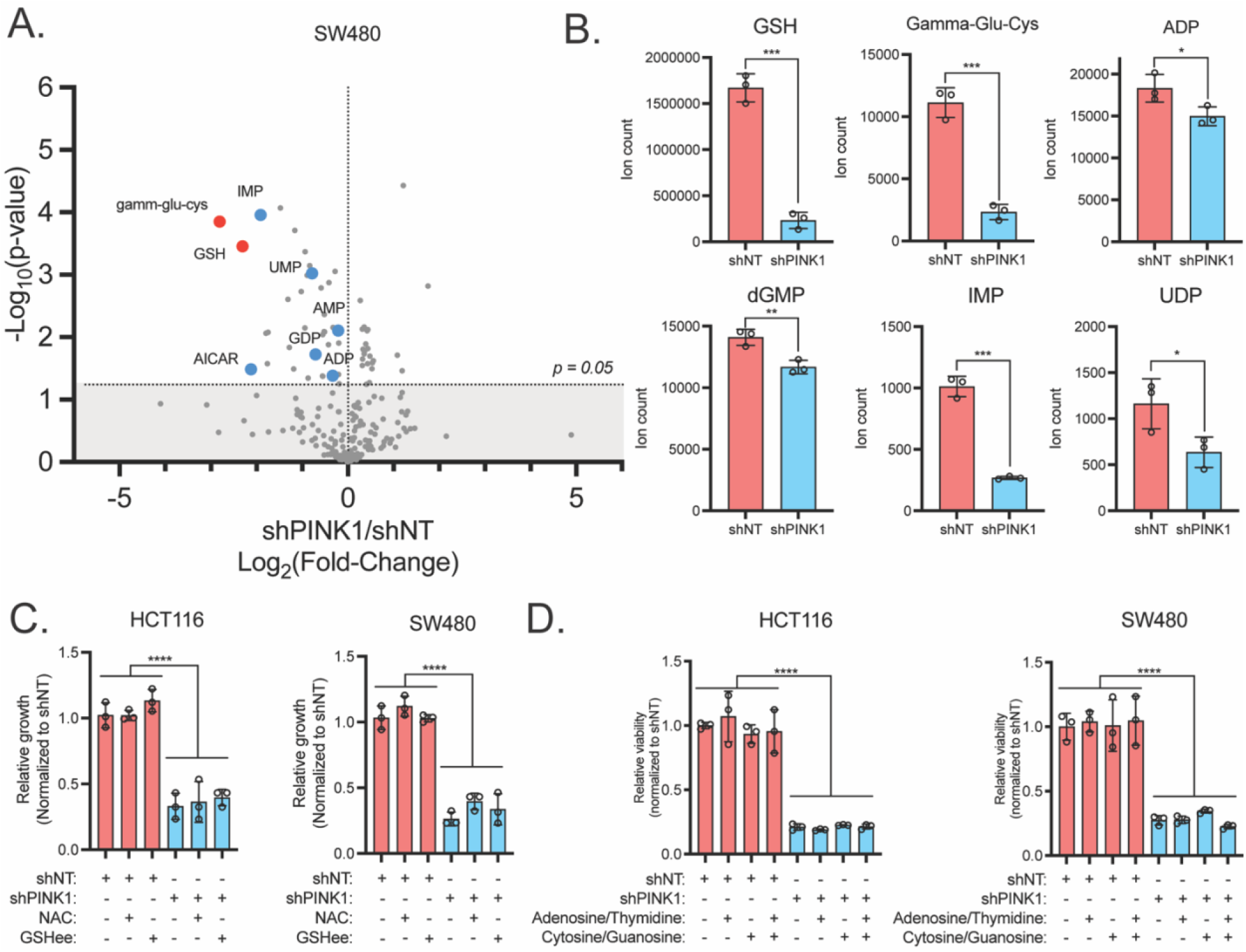
Disruption of redox and nucleotide metabolism is secondary to PINK1-dependent growth defects. **A**. Metabolite profiling via mass spectrometry-based metabolomics 3 days following induction of shNT and shPINK1 (n=3). **B**. Selected metabolites associated with redox and nucleotides from SW480 were shown to decrease under PINK1 knockdown. Cell growth following treatment with **C**.1mM NAC and GSHee or **D**. 100μM adenosine/thymidine or cytosine /guanosine. Data is presented as mean +/- the standard error of the mean, *P < 0.05, **P < 0.001, ****P < 0.0001. All experiments were done in triplicates and repeated at least three times, the metabolomics were done with N = 3 per condition.

To reverse proliferation defects that resulted from PINK1 KD, based on our metabolomics data, we supplemented cells with the antioxidants N-acetylcysteine (NAC) or glutathione ethyl ester (GSH-ee), or the nucleosides (adenosine, thymidine, cytosine, and guanosine) (**Figure 4C and 4D**). However, these metabolites did not rescue the growth defects in PINK1 KD cells. Since mitochondrial dysfunction can contribute to elevated NADH/NAD ratio, we assayed NADH/NAD ration in PINK1 KD cells. We observed elevated NADH/NAD ration upon PINK1 KD. To resolve elevated NADH/NAD ratio, we employed *Lactobacillus brevis* NADH oxidase (*Lb*NOX) and mitochondrial targeted LbNOX (mt*Lb*NOX), which oxidize NADH to water, in order to determine the impact of reductive stress on our PINK1 KD phenotype. Here too, we failed to rescue PINK1 KD^19^ (**Supplemental figure 2A**). Lastly, the pan-caspase/apoptosis inhibitor (z-VAD-FMK), ferroptosis inhibitor (Ferrostatin 1; Fer1), and necroptosis inhibitor (Necrostatin 1; Nec1) did not reverse growth defects in PINK1 KD cells (**Supplemental figure 2B**). Collectively, these data suggested that metabolic dysregulation was secondary of the growth suppressive and mitochondrial dysfunction phenotypes following PINK1 KD.

### PINK1 modulates intracellular iron distribution independent of the canonical iron starvation response

In addition to their above noted role in metabolism, mitochondria are also important hubs for cellular iron homeostasis. Iron sulfur clusters (Fe-S) and heme biosynthetic pathways initiate in the mitochondria. Iron and iron-containing cofactors are critical for electron transport and redox balance as iron is a redox active element that can shuttle electrons along the respiratory chain. Moreover, we have shown that nucleotide metabolism requires iron for pyrimidine/purine biosynthesis^20^. Acute depletion of mitochondrial iron via deferiprone (DFP) induces mitophagy, thus linking mitochondrial turnover to cellular iron homeostasis^11^. To measure the mitochondrial iron pool, we utilized a flow cytometry compatible mitochondrial iron stain, Mito-FerroGreen. With this, we observed that PINK1 KD decreased mitochondrial iron levels (**Figure 5A**).

**Figure 5.**
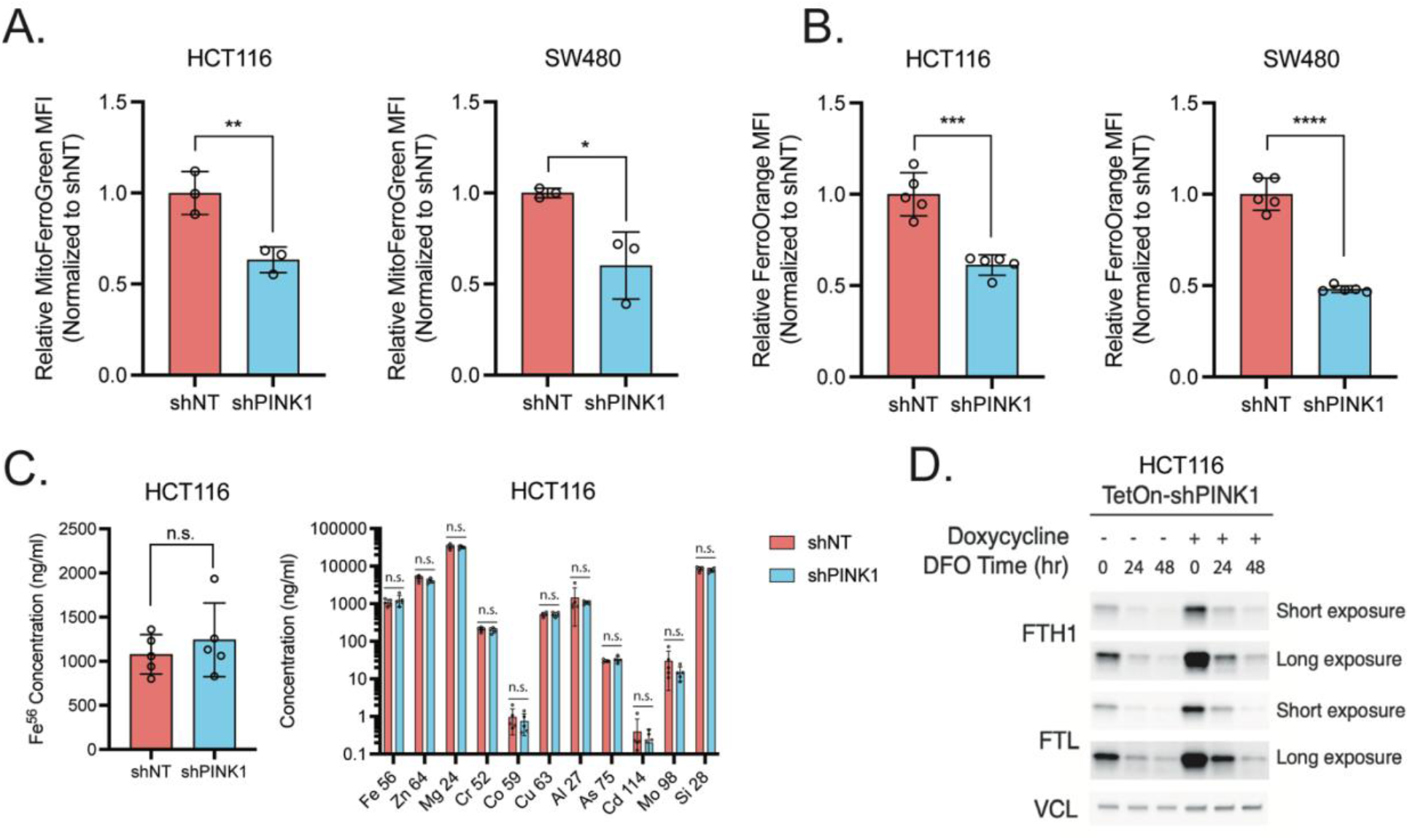
PINK1 modulates intracellular iron distribution independent of canonical iron starvation response. **A**. MitoFerroGreen and **B**. FerroOrange quantitation in shNT and shPINK1 cells by flow cytometry analyses of mean fluorescence intensity (MFI). **C**. ICP-MS analyses of divalent metals in shNT and shPINK1 cells. **D**. Immunoblotting of WT and shPINK1 cells treated with 100 μM of deferoxamine (DFO) for 24 and 48 hour. Data is presented as mean +/- the standard error of the mean, *P < 0.05, **P < 0.001, ****P < 0.0001. All experiments were done in triplicates and repeated at least three times, the ICP-MS were done with N = 5 per condition.

Mitochondrial iron is sensitive to cytosolic iron perturbation. For example, extracellular iron depletion induces HIF activation, and thus drives expression of mitochondrial iron transport SLC25A37^21,22^. Cells have an extensive regulatory network that monitors iron homeostasis^23^. Excess cytosolic labile iron pool (LIP) can lead to oxidative damage^24,25^. Thus, iron is tightly regulated through many overlapping and distinct pathways. Using FerroOrange to measure the LIP, we observed that this was robustly decreased upon PINK KD (**Figure 5B**). In contrast, total cellular iron was not changed, as assessed by inductively coupled plasma-mass spectrometry (ICP-MS), as well as other trace metal elements (**Figure 5C**). High levels of iron lead to the upregulation of the iron storage protein ferritin (FTN) expression and stabilization, which is composed of ferritin heavy chain (FTH1) and ferritin light chain (FTL). FTN sequesters excessive iron, preventing cellular oxidative damage from excessive LIP. In PINK1 KD cells, increased FTH1 and FTL was observed (**Figure 5D**).

### Restoring iron homeostasis via NCOA4-mediated ferritinophagy reverses growth suppression upon PINK1 inhibition

To understand the role of iron in the growth defects following PINK1 KD, we supplemented with ferric ammonium citrate (FAC) to rescue LIP and mitochondrial iron. Interestingly, growth was not rescued in the PINK1 KD cells (**Figure 6A**). An increase in FTN levels upon FAC treatment was observed, but no increase in LIP or mitochondrial iron, indicating that introduction of exogenous iron was primarily integrated to FTN complexes (**Figure 6B and 6C**). Indeed, iron chelation with DFO decreased LIP (**Figure 6D**). Cells compensate for the loss of cellular LIP via a cargo-specific autophagic degradation of FTN by ferritinophagy^26,27^. We show that deferoxamine (DFO) depleted FTN levels in WT cells, but PINK1 KD maintained higher levels FTN (**Figure 5D**). This data suggested that there are defects in ferritinophagy that decrease LIP in the PINK1 KD cells.

**Figure 6.**
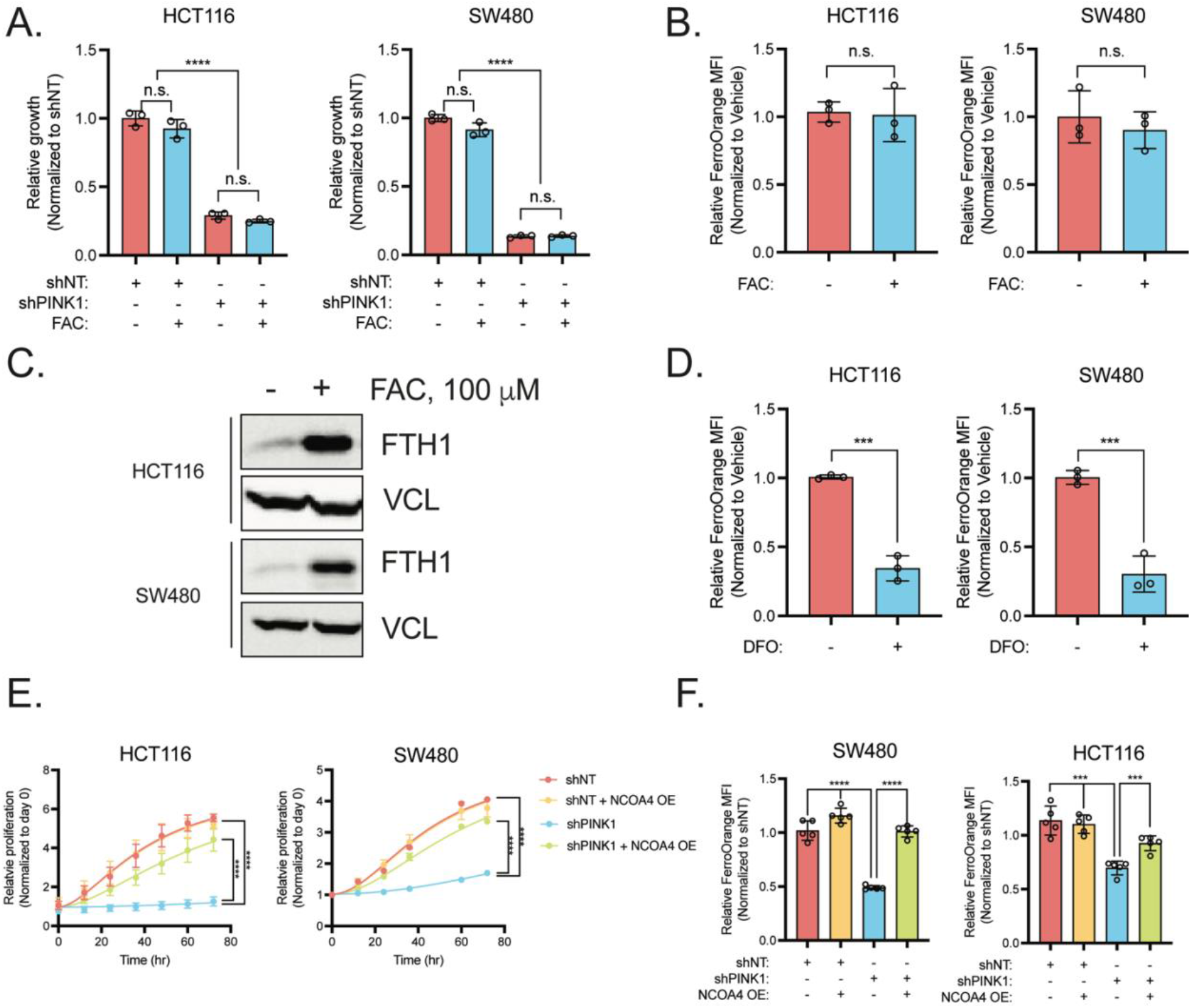
Restoring iron homeostasis via NCOA4-mediated ferritinophagy rescues growth in following loss of PINK1. **A**. shNT and shPINK1 cells were treated with 100 μM ferric ammonium citrate (FAC) and proliferation was measured for 72 hours. **B**. HCT116 and SW480 were treated with 100 μM FAC for 72 hours and the labile iron pool (LIP) was assessed by FerroOrange. **C**. Immunoblot of ferritin heavy chain (FTH1) following 100 μM FAC treatment. **D**. Measurement of LIP by FerroOrange upon DFO treatment. **E**. Cell growth or F. FerroOrange staining following NCOA4 overexpression in shNT and shPINK1 cells (mean fluorescence intensity (MFI)). Data is presented as mean +/- the standard error of the mean, *P < 0.05, **P < 0.001, ****P < 0.0001. All experiments were done in triplicates and repeated at least three times, the FerroOrange staining was done with N = 4 per condition.

Ferritinophagy is coordinated by the cargo receptor NCOA4, which binds to both FTL and FTH1 and delivers FTN to autophagosomes for degradation^27^. In addition to liberating FTN bound iron for other iron dependent processes, NCOA4 has been reported to be important for mitochondrial iron balance and respiration^28^. To rescue FTN accumulation in PINK1 KD cells, NCOA4 was over expressed (OE). NCOA4 expression was sufficient to rescue proliferative defects of PINK1 KD and restore labile iron pool (LIP) in these cells (**Figure 6E and 6F**). Overall, here we demonstrated that PINK1 loss leads to mitochondrial dysfunction and iron accumulation in ferritin. Liberating the sequestered iron from the ferritin complex by inducing ferritinophagy was able to compensate for PINK1 KD. This data suggests an essential role of mitophagy in regulating the LIP via ferritnophagy.

## Discussion

Mitochondria are important biosynthetic and metabolic hubs in tumorigenesis, and functional mitochondria are critical to support cancer cell growth^29^. Environmental stressors in the tumor microenvironment such as hypoxia and nutrient dysregulation and intrinsic factors driving mitochondrial DNA heteroplasmy require adaptations to mitochondrial dynamics, turnover, and programs for cancer cell survival ^30,31^. The induction of mitophagy requires exogenous expression of parkin and mitochondrial damage to remove the mitochondrial pool^32^. Mitophagy is essential in colon cancer growth, however the underlying mechanisms are unclear. The present work demonstrates that inhibition of PINK1-mediated mitophagy decreases colon cancer growth in a panel of colon cancer-derived cell lines. CRC cells have high basal mitophagy^16^ and failure to execute mitophagy leads to proliferative defects. Mechanistically, we show that colon cancer cells are dependent on PINK1 to maintain mitochondrial respiration. Although we did observe metabolite changes such as decreased reduced glutathione and several nucleotide species, simply added back these metabolites are not sufficient to restore PINK1 dependent proliferative defects. Rather, we demonstrate that PINK1 KD decreases the LIP and restoring ferritinophagy was sufficient to rescue cell proliferation.

The critical role of mitochondria in iron homeostasis is well appreciated. Mitochondrial uptake of iron via the mitoferrins (SLC25A27/38) is essential for the biosynthesis of iron sulfur clusters proteins and heme^33^. Iron sulfur clusters and heme are both important cofactors that support ETC, nucleotide biosynthesis, and DNA replication^34^. Although iron is a critical micronutrient essential for many biological processes, excessive free iron can be cytotoxic and exert oxidative stress.^35^ Therefore, the balance between iron storage in multimeric ferritin complexes and ferritin turnover via ferritinophagy is tightly regulated. In addition to its role in FTN turnover, the ferritinophagy adaptor NCOA4 has also been implicated in mitochondrial iron balance^12^. However, the canonical mitophagy pathway driven by PINK1, and how this cross-talks with cellular iron homeostatic mechanisms, are still unclear.

The most well studied PINK1 substrates include the mitochondrial outer membrane proteins mitofusins (MFN1/2), ubiquitin, and parkin^36^. The phosphorylation targets downstream of PINK1 activation are often signals that recruit autophagy adaptors to delivery mitochondria to autophagosomes. The kinase domain of PINK1 faces the cytosolic side and is postulated to have targets outside of OMM proteins^37^. Interestingly, phosphorylation targets for PINK1 were identified under mitochondrial uncoupled conditions. However, our study pinpoints a role of homeostatic PINK1 in CRC cells. This distinction may alter the subset of proteins targeted by PINK1 basally. Although mitophagy and ferritinophagy are distinct cargo-selective autophagic pathways, several players in these two pathways are shared^38^. We speculate that loss of PINK1 inhibits overlapping effectors that are central in mitophagy and ferritinophagy. Thus, future studies will be needed to identify relevant PINK1 targets and the complex interplay of various autophagy adaptors in balancing mitophagy and ferritinophagy.

We provide evidence of direct crosstalk between two cargo-specific autophagic pathways, mitophagy and ferritinophagy. This connection between mitochondrial quality control mechanism and iron homeostasis has been reported to mediate important cellular pathophysiology, but mechanistic insights have been lacking. A study suggested the possibility that NCOA4 recognizes mitochondrial ferritin (FTMT) on the outer mitochondria membrane and directly degrading mitochondria via autophagy^12^. However, we propose a potential alternative mechanism to their NCOA4-dependent mitophagy paradigm since FTMT expression is undetectable in our CRC cell lines. Previous studies have identified that intestinal epithelial derived cancer cells utilize mitophagy to evade T cell immunity via lysosomal iron overload^39^. In addition, genetic deletion of PINK1 in pancreatic cancer contributes to mitochondrial iron overload via accumulation of mitochondrial iron importers^21^. Since mitochondria can harbor abundant cellular iron, their roles in mediating iron storage and subcellular utilization have been increasingly scrutinized^40,41^. Moreover, colon cancer cells sequester iron to fuel pro-survival responses such as hypoxic signaling and nucleotide biosynthesis. Due to the redox reactivity of iron, iron mobilization is swiftly carried out by chaperones and compartmentalized by different organelles. Disruption of iron handling machineries have been implicated in cellular dysfunction. Loss of mitochondrial iron contributes to decreased oxygen consumption and respiratory chain defects.

Mitochondrial and ferritin turnover coordinate dynamic responses central in sustaining CRC growth. Although there are no specific agents that specifically block mitophagy, hydroxychloroquine is a lysosomotropic agent that blocks all form of autophagy consequent with a neutralization of lysosomal pH and lysosome dysfunction. Hydroxychloroquine has demonstrated promising results in conjunction with standard chemotherapeutics in metastatic CRC patients (NCT01206530) and our work suggests assessing the role of cellular iron availability in the efficacy of hydroxychloroquine in these patients. Moreover, recent work has identified compounds that directly inhibit ferritinophagy and may lead to more direct and potent growth inhibition of CRC^42^.

### Experimental procedures

#### Cell line and culture

HCT116, DLD1, MC38, HT29 and SW480 cells were maintained at 37°C in 5% CO2 and 21% O2. Cells were cultured in Dulbecco’s Modified Eagle Medium (DMEM) supplemented with 10% FBS and 1% antibiotic/antimycotic. Constructs for doxycycline inducible shRNA were generated using the Tet-pLKO-puro (Dmitri Wiederschain; Addgene plasmid #21915). shRNA primer sequences are as follows: shPRKN (F: CCGGGCTTAGACTGTTTCCACTTATCTCGAGATAAGTGGAAACAGTCTAAGCTTTTT, R: AATTAAAAAGCTTAGACTGTTTCCACTTATCTCGAGATAAGTGGAAACAGTCTAAGC), shPINK1.2 (F: CCGGGAAATCTTCGGGCTTGTCAATCTCGAGATTGACAAGCCCGAAGATTTCTTTTT, R: AATTAAAAAGAAATCTTCGGGCTTGTCAATCTCGAGATTGACAAGCCCGAAGATTTC), shPINK1.3 (F: CCGGGCCGCAAATGTGCTTCATCTACTCGAGTAGATGAAGCACATTTGCGGCTTTTT, R: AATTAAAAAGCCGCAAATGTGCTTCATCTACTCGAGTAGATGAAGCACATTTGCGGC). CRISPR knockout line was generated using gRNA in LenticrisprV2 (Feng Zhang; Addgene plasmid 49535). CRISPR primer sequences are as follows: sgBNIP3 (F: CACCGATGGGATTGGTCAAGTCGGC, R: AAACGCCGACTTGACCAATCCCATC), sgNIX/BNIP3L (F: CACCGCGGCGGCGGCTCGACTAGGT. R: AAACACC TAGTCGAGCCGCCGCCGC), sgFUNDC1 (F: CACCGTAATGGGTGGCGTTACTGGC, AA ACGCCAGTAACGCCACCCATTAC). NCOA4 overexpression plasmid (pIND20-NCOA4) was generously donated from Joseph D. Mancias’ group. Plasmids were generated and inserted in to a lenti-viral vector for stable transfection. Knockdown was induced using 500 ng/mL of doxycycline for 48-hours. Ferric ammonium citrate was obtained from Sigmal Aldrich (RES20400-A7). CCCP (25458), Deferoxamine mesylate (14595), Ferrostatin-1 (17729), Z-VAD(OH)-FMK (14467), Necrostatin-1 (11658), and doxycycline hyclate (14422) was obtained from Cayman Chemicals.

#### Mouse xenograft model

Immunocompromised (NOD.Cg-Prkdc^scid^/J), 6- to 8- or 8- to 10-week-old mice of both sexes were maintained in the facilities of NCI Frederick National Laboratory. For subcutaneous xenograft studies, HCT116 and SW480 or shNT, shPINK1.2 and shPINK.13 cells were trypsinized and 2 million cells were implanted into the lower flanks. All treatments began on day 10 after tumors became visible. 200mg dox/kg diet was used to induce shRNA expression. The diet was purchased from Bioserv. Subcutaneous tumor size was measured with digital calipers at the indicated time points. Tumor volume (V) was calculated as V = 1/2(length × width2). At the endpoint, mice were sacrificed and tumors were excised. The final tumor volume and weight were measured, and tissue was used for proliferation and apoptosis assay.

#### Proliferation assay

Growth assays were performed using MTT (Thiazolyl Blue Tetrazolium Bromide) assays and live cell imaging. Briefly, for MTT cells were plated down and 24-h following plating a Day 0 reading was taken. Cells were incubated for 45 min with MTT solution (5× concentrate stock: 5 mg/mL, in 1XPBS, pH 7.4). Media and MTT solution were then carefully aspirated followed by solubilization with dimethyl sulfoxide. Absorbance was read at 570nm. Following the Day 0 read, the cells were treated with indicated doses and readings were taken after 24-h. Live cell imaging was done using the Cytation 5 Imaging Multi-Mode reader. Cells were plated down, treated 24 h later with indicated treatments, and immediately imaged and analyzed for cell number. Images were then taken every 24 h.

#### Real time quantitative PCR

1 μg of total RNA extracted using Trizol reagent from mouse tissues human intestinal cell lines. RNA was reverse transcribed to cDNA using SuperScriptTM III First-Strand Synthesis System (Invitrogen). Real time PCR reactions were set up in three technical replicates for each sample. cDNA gene specific primers, SYBR green master mix was combined, and then run in QuantStudio 5 Real-Time PCR System (Applied BioSystems). The fold-change of the genes were calculated using the ΔΔCt method using β-actin as the housekeeping gene. Primers are listed as follows: human PINK1 (F: GCCTCATCGAGGAAAAACAGG. R: GTCTCGTGTCCAACGGGTC), mouse PINK1 (F: TTCTTCCGCCAGTCGGTAG. R: CTGCTTCTCCTCGATCAGCC), human PRKN/PARK2 (F: GTGTTTGTCAGGTTCAACTCCA. R: GAAAATCACACGCAACTGGTC), human NIX/BNIP3L (F: ATGTCGTCCCACCTAGTCGAG. R: TGAGGATGGTACGTGTTCCAG), human BNIP3 (F: CAGGGCTCCTGGGTAGAACT, R: CTACTCCGTCCAGACTCATGC), human FUNDC1 (F: CCTCCCCAAGACTATGAAAGTGA, R: AAACACTCGATTCCACCACTG), human beta-actin/ACTB (F: CATGTACGTTGCTATCCAGGC, R: CTCCTTAATGTCACGCACGAT), mouse beta-actin/ACTB (F: GTGACGTTGACATCCGTAAAGA, R: GCCGGACTCATCGTACTCC).

#### Western blotting

Whole-cell lysate preparations were described previously (Anderson et al., 2013). Whole cell lysates were prepared from cell lines by RIPA buffer. Homogenates were incubated in RIPA buffer for 15 min on ice followed by 13,000 rpm centrifugation for 15 min. Supernatants were transferred to a new tube and mixed with 5× Laemmli buffer and boiled for 5 min. Lysates containing 30–40 μg of protein per well were separated by SDS-PAGE, transferred onto nitrocellulose membranes, and immunoblotted overnight at 4°C with indicated antibodies: Phospho-Ubiquitin (Ser65) (E2J6T) Rabbit mAb #62802 (Cell Signaling Technology), Vinculin (E1E9V) XP Rabbit mAb #13901(Cell Signaling Technology), Anti-ferritin heavy chain Antibody (B-12): sc-376594 (Santa Cruz Biotechnology), Anti-ferritin light chain Antibody (D-9): sc-74513 (Santa Cruz Biotechnology). All the primary antibodies were used at a dilution of 1:1000. HRP-conjugated secondary antibodies used were anti-rabbit and anti-mouse at a dilution of 1: 2000 and immunoblots were developed using Chemidoc imaging system (ChemiDoc, BioRad).

#### NAD/NADH measurement

NAD/NADH-Glo assays from Promega (G9071) were purchased and performed based on manufacturer instructions.

#### Clonogenic assays

Cells were plated in six-well plates in biological triplicates at 300–600 cells per well in 2 mL of media. Dox-media were changed every 2 days. Assays were concluded after 10–15 days by fixing in –20 °C cold 100% methanol 10 min and staining with 0.5% crystal violet 20% methanol solution for 15 min. Colonies were quantified using ImageJ or manually counted.

#### FerroOrange and Mito-FerroGreen measurement

The cells were washed with HBSS three times. Mito-FerroGreen working solutions (5 μmol/l) and FerroOrange working solution (1 μmol/l) were added to the cells, and the cells were incubated at 37°C for 30 minutes in a 5% CO2 incubator. The supernatant was discarded and the cells were washed with HBSS three times. Cells were then processed for flow cytometry analyses.

#### Mitophagy assay with flow cytometry

Cells were seeded at 100,000 cells per well in a twelve-well plate. Next day, 500ng of pCLBW cox8 EGFP mCherry (David Chan;Addgene plasmid #78520).) were transfected into the cells with Lipofectamine 2000 according to manufacturer’s protocol. Cells were then treated with doxycycline for 48hour to induce PINK1 KD. After 48hour post transfection, cells were trypsinized and collected for flow cytometry analyses. Cells were counterstained with DAPI to gate out dead cells. Then, mCherry positive cells were gated and EGFP fluorescence was assessed. Ratio of low EGFP to high EGFP was plotted as % mitophagy. Analysis was done using FlowJo software.

#### Metabolomics

Cells were plated at 500,000 cells per well in six-well plates or ∼1.5 million cells per 10-cm dish. At the endpoint, cells were lysed with dry-ice cold 80% methanol and extracts were then centrifuged at 10,000 × g for 10 min at 4 °C and the supernatant was stored at –80 °C until further analyses. Protein concentration was determined by processing a parallel well/dish for each sample and used to normalize metabolite fractions across samples. Based on protein concentrations, aliquots of the supernatants were transferred to a fresh microcentrifuge tube and lyophilized using a SpeedVac concentrator. Dried metabolite pellets were re-suspended in 45 μL 50:50 methanol:water mixture for LC–MS analysis.

The QqQ data were pre-processed with Agilent MassHunter Workstation Quantitative Analysis Software (B0700). Each sample was normalized by the total intensity of all metabolites to scale for loading. Finally, each metabolite abundance level in each sample was divided by the median of all abundance levels across all samples for proper comparisons, statistical analyses, and visualizations among metabolites. The statistical significance test was done by a two-tailed t-test with a significance threshold level of 0.05.

#### Induced coupled plasma mass spectrometry (ICP-MS)

Metal quantifications by ICP-MS were performed as previously described^13^. Briefly, tissue samples were digested with 2 mL/g total weight nitric acid (BDH Aristar Ultra) for 24 h and then digested with 1 mL/g total weight hydrogen peroxide (BDH Aristar Ultra) for 24 h at room temperature. Specimens were preserved at 4ºC until quantification of metals. Ultrapure water was used for the final sample dilution. Samples were analyzed using a Perkin-Elmer Nexion 2000 ICP-MS.

#### Seahorse mito stress test

Cells were seeded at 2 × 104 cells/well in 80 μl/well of normal growth media (DMEM with 25 mM Glucose and 2 mM Glutamine) in an Agilent XF96 V3 PS Cell Culture Microplate (#101085-004). To achieve an even distribution of cells within wells, plates were incubated on the bench top at room temperature for 1 h before incubating at 37 °C, 5% CO2 overnight. To hydrate the XF96 FluxPak (#102416-100), 200 μL/well of sterile water was added and the entire cartridge was incubated at 37 °C, no CO2 overnight. The following day, 1 h prior to running the assay, 60 μL/well of growth media was removed from the cell culture plate, and cells were washed twice with 200 μL/well of assay medium (XF DMEM Base Medium, pH 7.4 (#103575-100) containing 25 mM glucose (#103577-100) and 2 mM glutamine (#103579-100)). After washing, 160 μL/well of assay medium was added to the cell culture plate for a final volume of 180 μL/well. Cells were then incubated at 37 °C, without CO2 until analysis. One hour prior to the assay, water from the FluxPak hydration was exchanged for 200 μL/well of XF Calibrant (#100840-000) and the cartridge was returned at 37 °C, without CO2 until analysis. Oligomycin (100 μM), FCCP (100 μM), and Rotenone/Antimycin (50 μM) from the XF Cell Mito Stress Test Kit (#103015-100) were re-constituted in assay medium to make the indicated stock concentrations. Twenty microliters of Oligomycin was loaded into Port A for each well of the FluxPak, 22 μL of FCCP into Port B, and 25 μL of Rotenone/Antimycin into Port C. Port D was left empty. The final FCCP concentration was optimized to achieve maximal respiration in each condition.

The Mito Stress Test was conducted on an XF96 Extracellular Flux Analyzer and OCR was analyzed using Wave 2.6 software. Following the assay, OCR was normalized to cell number with the CyQUANT NF Cell Proliferation Assay (C35006) from Thermo Fisher according to manufacturer’s instructions.

#### Quantification and statistical analysis

In vitro experiments were validated in 4 cell lines. Each cell line experiment was performed in technical replicates for each condition and repeated at least three times with biological triplicates to ensure reproducibility. Figures show a representative biological replicate unless otherwise indicated. Blinding was performed whenever appropriate. Sample description and identification was unavailable to the core personnel during data collection and analysis. Statistical details of all experiments can be found in the figure legends. The sample numbers are mentioned in each figure legend and denote biological replicates. Statistical details are reported in figure legends. Results are expressed as the mean plus or minus the standard error of the mean for all figures unless otherwise noted. Significance between 2 groups was tested using a 2 tailed unpaired t test. Significance among multiple groups was tested using a one-way ANOVA. GraphPad Prism 7.0 was used for the statistical analysis. Statistical significance is described in the figure legends as: * p < 0.05, ** p < 0.01, *** p < 0.001, **** p < 0.0001.

## Acknowledgements

This work was funded by NIH grants: R01CA148828, R01CA245546, and R01DK095201 (Y.M.S); R37CA237421, R01CA248160, and R01CA244931 (C.A.L); R01 DK124384 (J.D.M); UMCCC Core Grant P30CA046592 and Center for Gastrointestinal Research P30DK034933 (Y.M.S and C.A.L). B.C. was supported by a Department of Defense National Defense Science and Engineering Graduate Fellowship (DoD NDSEG) and NIH Cellular and Molecular Biology Training Grant T32-GM145470.

