## Supplemental Figures 1 and 2 for "PINK1 supports colorectal cancer growth by regulating the labile iron pool"

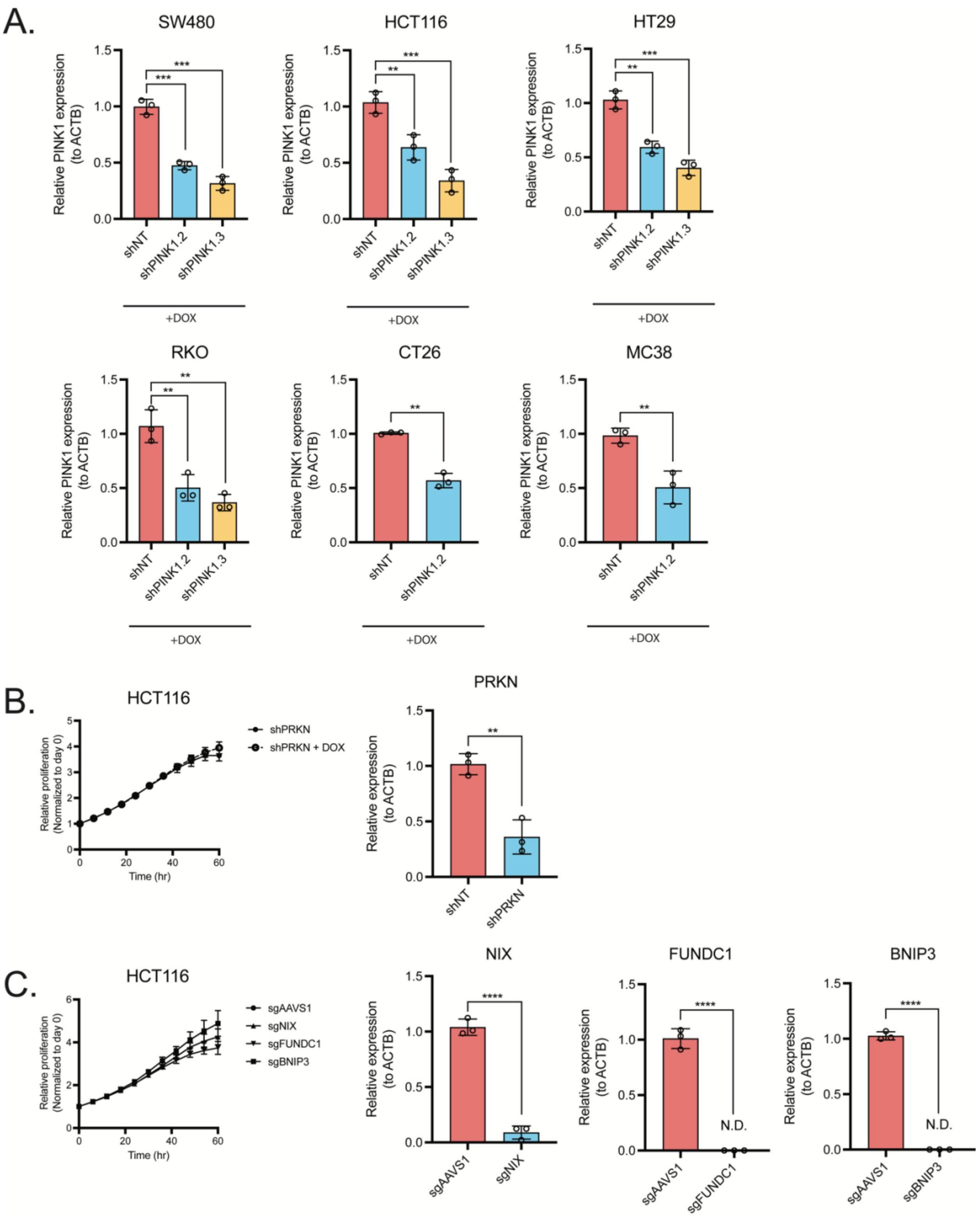


**Supplemental figure 1. PINK1 is required for colorectal cancer in vitro and in vivo growth, but not Parkin or other mitophagy receptors.** A. qRT-PCR data of PINK1 mRNA transcript level in shNT, shPINK1.2, and shPINK1.3 in SW480, HCT116, HT29, CT26, and MC38. B. HCT116 growth and qRT-PCR data of shRNA targeting PAKN. C. HCT116 growth and qRT-PCR data of sgRNA targeting control (sgAAVS1), and mitophagy receptors sgNIX, sgFUNDC1, and sgBNIP3. N.D. denotes non-detectable. (mean -/+ SEM, N = 3 per condition).** indicates P < 0.001, **** indicates P < 0.0001.

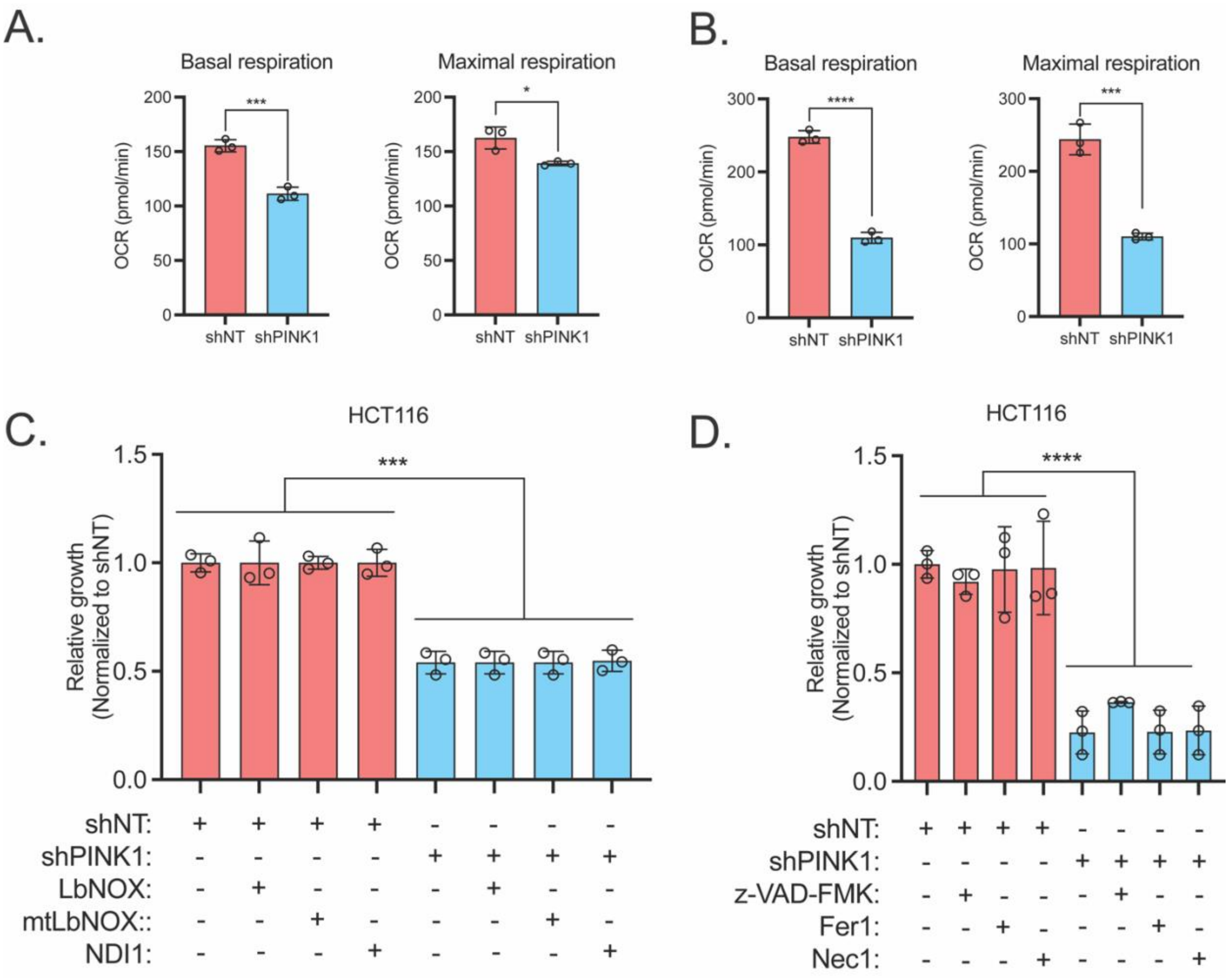


**Supplemental figure 2. Restoring NADH/NAD ratio, electron transport chain (ETC) complex I (CI), and cell death do not rescue PINK1 KD.** Basal and maximal respiration of A. SW480 and B. HCT116. Relative growth of shNT and shPINK1 cells C.expressing LbNOX, mtLbNOX, and NDI1, and treated with D. apoptosis inhibitor z-VAD-FMK (5 µM), ferroptosis inhibitor Ferostatin 1 (Fer1) (1 µM), and necroptosis inhibitor Necrostatin 1 (Nec1) (1 µM). *** indicates $P < 0.001$, **** indicates $P < 0.0001$.
